# Kcnq2/Kv7.2 controls the threshold and bihemispheric symmetry of cortical spreading depolarization

**DOI:** 10.1101/2020.11.27.401570

**Authors:** Isamu Aiba, Jeffrey L Noebels

**Affiliations:** Department of Neurology, Baylor College of Medicine, Houston TX 77030

**Keywords:** epilepsy, seizure, Kv1.1, retigabine, XE991

## Abstract

Spreading depolarization (SD) is a slowly propagating wave of massive cellular depolarization associated with acute brain injury and migraine aura. Genetic studies link molecular defects in enhanced cytoplasmic Ca^2+^ flux, glial Na^+^-K^+^ ATPase, and interneuronal Na^+^ current with SD susceptibility, emphasizing the important roles of synaptic activity and extracellular ionic homeostasis in determining SD threshold. In contrast, gene mutations in ion channels that shape intrinsic membrane excitability are frequently associated with epilepsy susceptibility. It is not well known whether epileptogenic mutations in voltage-gated potassium channels that regulate membrane repolarization also modify SD threshold and generation pattern. Here we report that the Kcnq2/Kv7.2 potassium channel subunit, frequently mutated in developmental epilepsy, is an SD modulatory gene with significant control over seizure-SD transition threshold, bilateral cortical expression, and temporal susceptibility. Chronic DC-band cortical EEG recording from awake conditional Kcnq2 deletion mice (Emx2^cre/+^::Kcnq2^flox/flox^) revealed spontaneous cortical seizures and SD. In contrast to a related potassium channel deficient model, Kv1.1 KO mice, spontaneous cortical SDs in Kcnq2 cKO mice are tightly coupled to the terminal phase of seizures, arise bilaterally, and are observed predominantly during the dark phase. Administration of the nonselective Kv7.2 inhibitor XE991 to Kv1.1 KO mice reproduced the Kcnq2 cKO-like SD phenotype (tight seizure coupling and bilateral symmetry) in these mice, indicating that Kv7.2 currents directly and actively modulate SD properties. *In vitro* brain slice studies confirmed that Kcnq2/Kv7.2 depletion or pharmacological inhibition intrinsically lowers the cortical SD threshold, whereas pharmacological activators elevate the threshold to multiple depolarizing and hypometabolic SD triggers. Together these results identify Kcnq2/Kv7.2 as a distinctive SD regulatory gene, and point to SD as a potentially significant pathophysiological component of KCNQ2-linked epileptic encephalopathy syndromes. Our results also implicate KCNQ2/Kv7.2 channel activation as an adjunctive therapeutic target to inhibit SD incidence.

## Introduction

Spreading depolarization (SD) is a slow-moving wave of cellular depolarization in the brain gray matter associated with human neurological deficits in the setting of acute traumatic brain injury, ischemic stroke, and migraine with aura ^1,2^. Recent experimental studies also implicate SD in sudden unexpected death in epilepsy (SUDEP ^3^) and early epileptogenesis in glioblastoma ^4^. SD begins as a localized severe depolarization of brain tissue that profoundly elevates the extracellular concentration of excitatory solutes such as potassium and glutamate which defuse and further depolarize surrounding tissue, creating a self-regenerative, spreading wave of cellular depolarization and edema ^1^. Cellular depolarization during SD is so severe that transmembrane ionic gradients collapse, transiently silencing the activity of affected networks or causing permanent damage when brain tissue is metabolically compromised. SD also promotes malignant neurovascular inflammation and may contribute to longer lasting neurological deficits ^5,6^.

Seizures and SD are related excitatory events and can be generated alone or in a temporally associated manner in isolated brain slices, intact and injured brains, and computational simulations ^7–11^. Cortical seizures elevate extracellular K^+^ concentration to a ceiling level of 7-12 mM ^12,13^, a concentration range close to the SD threshold ^1,14^, sometimes leading to secondary generation of SD ^8,15,16^. SD arising secondary to a seizure may contribute to postictal complications such as migraine and hemiplegia (e.g. Todd’s paralysis ^17^). However, not all seizures are associated with SD, and epilepsy patients do not always display postictal SD-related deficits. While their relationship is complex, no molecular determinants for this seizure-SD coupling threshold have yet been identified.

The growing identification of genes causally assigned to clinically defined epilepsy syndromes is providing decisive mechanistic insight into the molecularly diverse pathogenesis of seizure disorders. Similarly, while fewer in number, genetic analysis of complicated migraine with aura syndromes such as familial hemiplegic migraine (FHM) reveal molecular mechanisms underlying SD threshold and susceptibility. The genes identified in FHM so far share a risk of developmental epileptic co-morbidity via distinct mechanisms, including calcium current facilitation of synaptic glutamate release (FHM1: CACNA1A ^18,19^), impaired extracellular K^+^ clearance (FHM2: ATP1A2 ^20,21^), and defective inhibitory neuron excitability (FHM 3: SCN1A ^22,23^). These genes assign an SD threshold-determining role to excitatory /inhibitory synaptic function and ionic homeostasis in the interstitial space. In epilepsy, mutations in genes regulating intrinsic membrane excitability, including SCN1A, are a common cause of epilepsy with a wide range of phenotypic severity. Given the distant relationship between seizure and SD, genes among this group may also contribute to SD threshold, however it is not known whether they generate distinctive SD phenotypes.

To further explore the influence of intrinsic membrane excitability defects, we evaluated the ability of Kcnq2/Kv7.2 channel activity to affect cortical SD using a viable conditional Kcnq2 KO mouse model (Emx1-cre::Kcnq2^flox/flox^) and synthetic Kcnq2 channel modulators. Kcnq2 encodes the potassium channel subunit Kv7.2 that forms heterotetramers in axon membranes and generates non-inactivating hyperpolarizing M-type K^+^ current to set the resting membrane potential^24,25^. Mutations in human KCNQ2 are associated with a phenotypic spectrum of neonatal-onset epilepsies extending from benign neonatal to severe developmental encephalopathy ^26(p2),27–29^. We combined *in vivo*/*vitro* electrophysiology and Kcnq2 pharmacology to examine SD characteristics of Kcnq2 deficient mice and compare them with a second axonal potassium channel seizure model, Kv1.1 deficient mice. Our recordings from Kv1.1 KO and Kcnq2-cKO mice identified significant differences in their cortical SD phenotypes as well as a direct interdependence of these two axonal potassium channels on SD threshold.

## Materials and methods

### Animal

All animal experimental protocols were approved by Baylor College of Medicine IACUC. Experiments were conducted on WT C57BL6J, Emx1-cre::Kcnq2 floxed conditional KO (Kcnq2-cKO) mouse lines. Kcnq2-flox mice were a gift of Dr A. Tzingounis ^30^. Emx1-IRES-cre (JAX Stock No: 005628) mice were purchased from the Jackson Laboratory. Kv1.1 KO mice (Tac:N:NIHS-BC) were obtained from the BCM breeding colony (also available from Jackson laboratory (JAX Stock No: 003532). All animals were housed and bred in the Center of Comparative Medicine at Baylor College of Medicine.

### Study design and Statistics

*In vivo* DC monitoring sessions were carried out to capture at least 20 spontaneous SD events. In total, 16 Kcnq2 and 8 Kv1.1 KO and WT mouse pairs were implanted, while four of Kcnq2-cKO mice were excluded because of poor postsurgical recovery or visible damage at the implant site. The identity of the mouse genotype was not blinded since it could be behaviorally identified during careful monitoring, however the EEG data were reviewed by two experienced investigators.

*In vitro* acute brain slice experiments were carried out in a blinded manner by masking the identity of the tissue genotype, and due to this experimental design, N is not always equal between groups. In order to reduce the number of animals and increase statistical power, most *in vitro* drug experiments involved repeated measurements. Because of the robust pharmacological and genetic effects, we set a minimum N=6 for independent comparisons and N=4 for repeated measurements in the same slice. *In vivo* anesthetized mouse experiments were repeatedly performed in each mouse to increase the statistical power. In these studies, latency differences between the onset of the 1^st^ (baseline) and 2^nd^ (drug) evoked SD were compared. All statistical comparisons were made using an unpaired t-test for 2-group comparisons and repeated measures ANOVA with post hoc Tukey’s multiple comparisons test where appropriate unless mentioned in the text. Statistics were computed using Graphpad Prism (GraphPad) and R software. All data are presented as mean standard ± deviation. Imaging data were analyzed using Image-J software.

### Acute brain slice preparation

Mice were deeply anesthetized with i.p. avertin (bromoethanol) followed by intracardiac perfusion with 10 ml of dissection solution (110 mM NMDG, 8 mM MgSO_4_, 25 mM NaHCO_3_, 1.25 mM Na_2_HPO_4_, 3mM KCl, 10 mM glucose, 0.4 mM ascorbate, saturated with 95% O_2_/5% CO_2_ gas), and then decapitated. The brain was extracted into ice-cold dissection solution, sagitally hemisected, and 300 µm coronal slices were cut on a vibratome (Leica 1300s) and the ventral portion was trimmed. Slices were incubated in dissection solution for 5 minutes at 33°C, then transferred to a submerged slice chamber and maintained in ACSF (in mM: 130 mM NaCl, 3 mM KCl, 25 mM NaHCO_3_, 1.25 mM Na_2_HPO_4_, 10 mM glucose, 0.4 mM ascorbate, saturated with 95% O_2_/5% CO_2_ gas) at room temperature. Recurrent SD generation was difficult to reproducibly trigger in a WT cortical slice bathed in normal ACSF. Therefore, in order to facilitate SD generation with recoverability, we elevated the KCl concentration of ACSF from 3 mM to 5 mM throughout the experiments, and recordings were made at 32-33°C.

### Models of SD generation and detection *in vitro* acute brain slices

In separate experiments, cortical SD was induced by potassium chloride (KCl) bath application, KCl microinjection, Mg^2+^ free bath solution, and exposure to oxygen-glucose deprivation (OGD) solution. In all 3 models, SD’s were generated and detected using a glass electrode (1-2 MΩ, filled with ACSF) placed in cortical layer 2-3 (50 µm depth) while simultaneously monitoring the intrinsic optical signal changes (i.e. light transparency increase due to tissue swelling) using a CCD camera (DMK27BU, Imaging Source, acquisition rate: 0.2-0.5Hz). SD onset was determined by detecting the IOS signal coincident with the electrophysiological depolarization within the imaging field. SD propagation rate was calculated by tracking the propagating wavefront over time. Imaged data of SD propagation were converted to the ratio (ΔI/I_0_). Since a DC shift was not always detected if the SD faded before reaching the recording electrode, most SD latency and propagation parameters were based on IOS signals.

Bath KCl application was conducted by incrementally increasing the bath KCl concentration. Pairs of cortical slices (WT and cKO) were incubated in a chamber and monitored as they were serially exposed to ACSF containing elevated KCl (10 to 14 mM, 1 mM increment, 5 minutes each). The KCl concentration that triggered SD was considered as the threshold.

In some experiments, KCl microinjection was used to trigger SD to characterize its propagation pattern in physiological KCl concentration. A single injection of 1 M KCl was pressure ejected via a micropipette placed in Layer 2/3 using a picospritzer (pressure: 40 psi, pulse duration: 100-500 ms, estimated ejection volume: ∼20 nl). In other slices, SD was triggered by exposure to nominally Mg^2+^ free ACSF in which MgSO_4_ was eliminated in the ACSF. In order to enhance tissue excitability, the bath divalent cation concentration was not compensated.

Finally, some slices were challenged metabolically using oxygen/glucose deprivation (OGD). OGD-SDs were induced by superfusing the cortical slices in the ACSF equilibrated with 95%N_2_/5%CO_2_ and reduced glucose concentration (0–5 mM glucose concentration osmotically balanced to 10 mM with sucrose). For pharmacological tests, slices were preincubated for 5-10 minutes at 34^°^C in ACSF containing the test compound, and then further exposed to OGD solution containing the same test compound.

### *In vivo* cortical SD activity in anesthetized mice

For acute (non-survival) *in vivo* studies, mice were anesthetized by intraperitoneal injection of urethane (1.5 mg/kg). A craniotomy (1.0 mm diameter) was made over the somatosensory cortex and covered with a gelfoam pledget soaked with saline. A thinned skull region was created 2 mm anterior to the cranial window and ∼0.1mm silver wire was placed for electrophysiological DC recording. In order to reduce the variability of tissue oxygenation, a flow of O_2_ was supplied to the nares. All mice showed a spO2 >95% as measured by pulse oximetry on the hindlimb (MouseOX, Starr Life Sciences Corp, PA). In order to avoid the initial hyperglycemia spike evoked by anesthesia, SD experiments were conducted >30 minutes after anesthesia induction. *In vivo* cortical SD threshold was determined by sequential application of KCl (125, 250, 500, 1000 mM, 2 µl) directly applied to the pial surface within the cranial window as described previously ^16^. SD was detected electrophysiologically by a strong negative DC shift and an IOS signal (>640 nm light scatter) recorded with a CCD camera (DMK27BU, Imaging Source).

### Chronic cortical SD monitoring in freely behaving mice

Spontaneous unfiltered cortical EEG activity was recorded with chronic electrodes implanted in freely moving Kcnq2-cKO (Emx1^+/-^-kcnq2^flox/flox^), and WT mice (Emx1^WT/WT^-kcnq2^flox/flox^). Juvenile mice (P18-20, 12-18 g body weight) were anesthetized with isofluorane (3% induction, 1.5-2.0% maintenance), scalp hair removed, and the skin cleansed with iodine/70% ethanol, locally anesthetized by lidocaine/bupivacaine mix, and a midline incision was made to expose the skull. Four bilaterally symmetric burr holes were made (±2 mm lateral, ±1.5mm anterior/posterior) over the frontal and parietal cerebral surface, and 2 burr holes for ground electrodes over the cerebellum (±1 mm lateral, +1 mm posterior from lambda). Silicon-coated silver wires (0.5 mm diameter) were inserted into the subdural space. A microconnector connected to the leads was glued onto the skull and cemented using Metabond (Parkell Inc). After recovery, mice were injected with meloxicam (5 mg/kg, s.c.) for 3 days for postoperative analgesia. Recordings were started as early as 5 days after surgery. Mice were transferred to a transparent box and the implanted microconnectors were tethered to the DC amplifier (ADI bioamp). DC activity was chronically monitored using ADI powerlab software system. The recording room was maintained on a 12-hour dark-light cycle. Mice were able to freely explore and had access to water and food. A seizure was identified as a slow stereotypic sharp wave oscillation overriding a small DC shift (∼0.2 mV) while SD was detected as a negative DC shift >5 mV in peak amplitude and the depolarization lasting for at least 30 seconds.

### Drugs

Retigabine was obtained from Tocris and ML213 from Sigma. Both drugs were freshly dissolved in DMSO at 100 mM on the day of the experiment. XE991 was purchased from Alomone labs and Sigma.

### Data availability

Data supporting the findings of this study are available within the article.

## Results

### Spontaneous cortical SD in awake kcnq2-cKO mice

We first characterized spontaneous SD generation in Kcnq2-deficient mice using chronic DC-band EEG recordings in awake animals implanted with cortical surface EEG electrodes. Because homozygous Kcnq2 KO results in neonatal death (at P0-1) likely due to respiratory failure ^31^, we studied conditional Kcnq2 KO (Emx1-cre::Kcnq2^flox/flox^, hereafter Kcnq2-cKO) mice in which Kcnq2 is selectively deleted in forebrain excitatory cells as well as peripheral ganglia ^32^, allowing prolonged survival into adulthood. In order to capture the native early developmental epilepsy phenotype while minimizing post-surgical mortality, the bilateral cortical EEG electrodes were implanted on a postnatal day 19-22 mice (**Figure 1A**) and recordings were conducted >5 days after surgical recovery.

**Figure 1.**
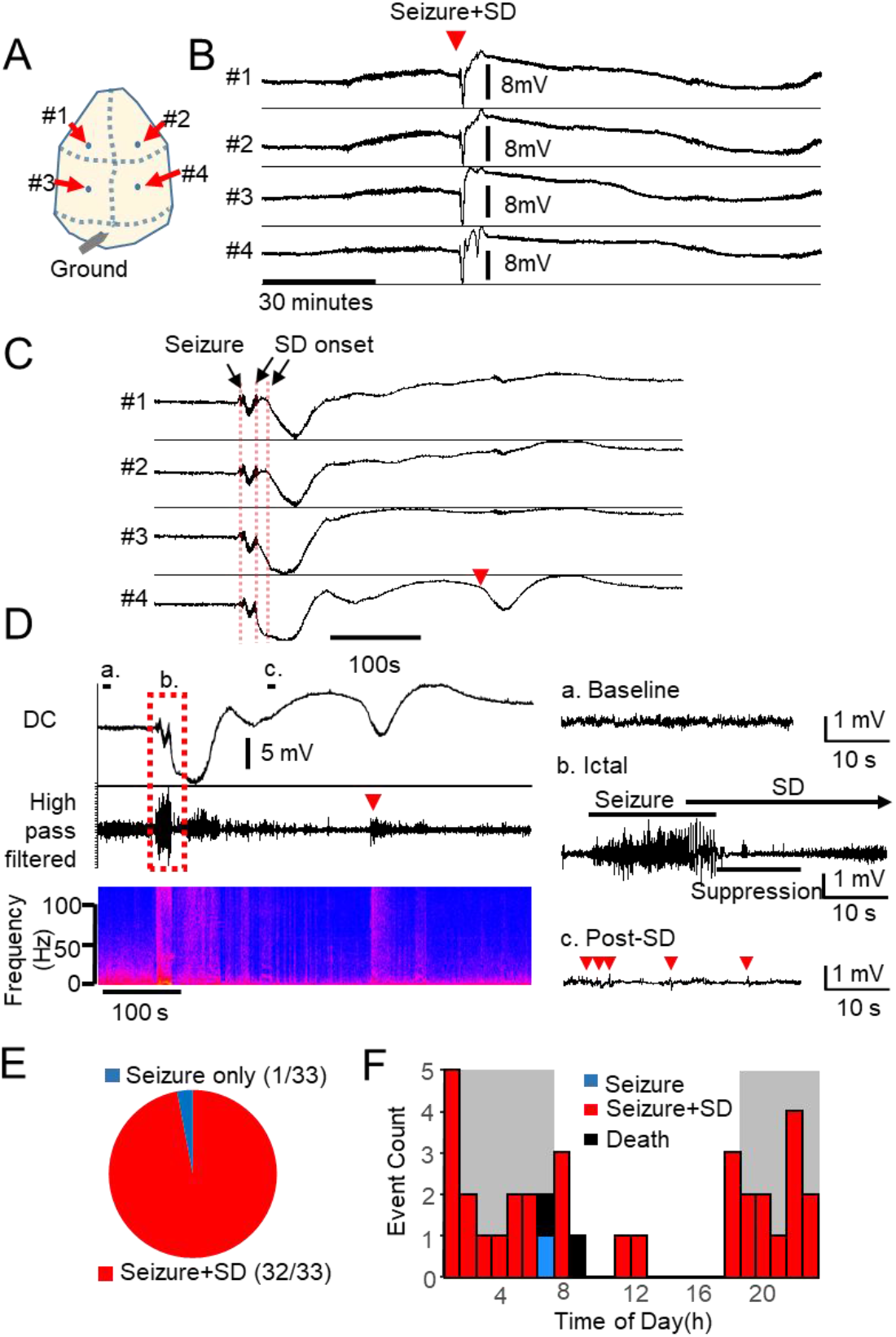
Spontaneous Seizure/SD complexes predominate in Emx1-Kcnq2cKO mice Chronic cortical DC recording from juvenile Kcnq2-cKO mouse. A. Recording configuration. B&C. Spontaneous seizure/SD complex in awake Kcnq2-cKO mouse shown at slow (B) and fast (C) time scales. SD was detected as a sharp negative DC potential shift following a brief generalized seizure detected simultaneously in both hemispheres. A solitary ectopic DC shift was occasionally detected posteriorly (C, arrowhead). D. characterization of fast EEG activity during seizure and SD. The negative DC shifts of SD were detected immediately after or even before the end of seizure activity. Left: slow time scale, right: faster scale. High-frequency EEG activity was enhanced during the nadir of the DC shift (b), followed by suppression with occasional spikelets (c). E. Almost all (97%, n= 34/35) spontaneous seizures were followed by SD. No SD’s were detected without a preceding seizure. F. Circadian cycle of seizure/SD complexes in 7 Kcnq2-cKO mice. Histogram of seizure and SD incidence during 24 h period (bin size = 1 hour, dark gray = dark phase). Spontaneous seizure/SD complexes were detected primarily during the dark phase (p<0.0001, Fisher’s exact test).

By optimizing the recording conditions, we obtained stable DC-band EEG recordings in freely behaving mice sufficient to detect sub-mV slow DC potential changes over multiple days of monitoring. During the chronic recording, we identified stereotyped large negative DC shifts, suggestive of SD, in association with generalized cortical seizure activity (**Figure 1B&C**). During a total of >1500 hours of chronic recordings from six Kcnq2-cKO mice, we detected 35 discrete SDs in the aftermath of seizure activity (0.6±0.4 SD per day, n=8 mice, total 67 days monitoring). The spontaneous seizure and SD events were tightly coupled; 97% (34/35) of seizures were followed by SD, and all SDs were preceded by seizure activity (**Figure 1E**). The SD generation pattern was regionally stereotyped, in that SDs were always first detected in the posterior cortex and were bilaterally synchronous (left vs right offset: 2.5 ± 3.6 s). The mean duration of seizure events was 23.4 ± 8.7 s, and the duration of the DC shift was 118 ± 51 s (anterior) and 100 ± 40 s (posterior). The mean SD amplitude was larger in posterior (13.3 ± 3.9 mV) than anterior (8.4 ± 3.1 mV) electrodes (p<0.001). SD propagation velocity (calculated based on the distance between posterior and anterior electrode pairs) was 8.4 ± 10.9 mm/minute. This high velocity is consistent with the fast SD propagation rate in Kcnq2-cKO cortical slices (discussed later in **Figure 4E&F**), however, spatial variability in SD generation/propagation pattern (e.g. generation between electrodes, see Discussion) may also contribute to the calculated velocity.

Following recovery from the large bilateral negative shift, unilateral ectopic slow DC shifts were occasionally detected at a posterior electrode (**Figure 1C**, red arrow), possibly representing a new isolated SD event or re-detection of a wandering SD. Thus, unlike the stereotypic SD generation (i.e. bilateral, posterior origin), the propagation pattern of each SD was somewhat variable. Two of the recorded mice died suddenly following a seizure-induced terminal SD with hind-limb extension, suggestive of SUDEP.

The spectral frequencies of fast cortical EEG activity during SD was analyzed in high-pass filtered (>0.1Hz) traces (**Figure 1D**). Electrographic seizures were characterized by synchronous high amplitude oscillatory activity (**Figure 1D, b. Ictal**), followed by a second period of brief EEG suppression during the negative slope of the DC shift. During the nadir of the negative DC shift, fast activity was enhanced, but after recovery from the negative DC shift, EEG activity was attenuated, while small spike and burst activities were detected (**Figure 1D, c. Post-SD**). The ectopic DC shift reactivated the EEG activity even during the post-SD EEG suppression (**Figure 1D left**, arrowhead). Video monitoring demonstrated that the initial seizure was convulsive, followed by post-ictal immobility during the SD phase. Overall, a single seizure led to a complex EEG and behavior abnormality sequence lasting 1-3 minutes in these animals.

No seizure or SD was ever detected in paired littermate WT control mice. In prolonged circadian recordings, we also found that 80% (28/35) of seizure-SD events in the mutants were detected during the dark phase, suggesting an influence of diurnal rhythm (**Figure 1F**). Together, chronic DC recordings revealed a unique aspect of the epileptic phenotype of Kcnq2-cKO mice characterized by tight seizure-SD coupling, bilateral SD generation/propagation, and a strong diurnal expression.

### Spontaneous seizure and SD in Kv1.1 KO mice

In order to compare the role of epilepsy-related potassium currents in SD, we made similar chronic recordings from age-matched juvenile Kcna1 KO mice. *Kcna1* encodes Kv1.1, a tetrameric voltage-gated potassium channel mediating delayed rectifier current widely expressed in the brain. Similar to Kcnq2, Kv1.1 is enriched within the axonal initial segment ^33,34^, and loss of these channels results in severe epilepsy in fly, mice, and humans ^35,36,37(p1),38,39^. The seizure phenotype in Kv1.1 KO mice is well documented, however *in vivo* cortical SD susceptibility in this model has not been studied.

In chronic recordings from eight Kv1.1 KO mice (P30-40), we identified 166 seizures (3.8 ± 5.0 per day) and 23 SD episodes (0.3 ± 0.2 per day) over 43 days of recording. Unlike Kcnq2-cKO mice, the co-occurrence of seizure and SD was far less common; of 166 seizures detected, only 4.8% (9/166) were associated with SD (PostIctal SD), and 14 SD episodes were detected without any preceding seizure activity (isolated SD, **Figure 2A&B, F**). The mean seizure duration was 52.1 ± 29.2 s, which is significantly longer than those detected in Kcnq2-cKO mice (Kcnq2-cKO: n=32, Kv1.1 KO: n=166, p<0.0001, Mann Whitney U-test). In Kv1.1 ko mice, the mean durations of SD were 97.52 ± 44.6 s (anterior) and 82.17 ± 27.0 s (posterior), and the amplitudes were 8.9 ± 5.4 mV (anterior) and 7.2 ± 5.3 mV (posterior). The calculated mean SD propagation velocity was 3.2 ± 2.5 mm/min, which is much slower than in Kcnq2-cKO (p<0.001, Mann Whitney U-test). Remarkably, all 9 postictal SDs were generated following a complete termination of seizure activity with a variable mean latency (24 ± 24 s), in contrast to Kcnq2-cKO mice, in which SDs were generated before full termination of the ongoing seizure activity.

**Figure 2.**
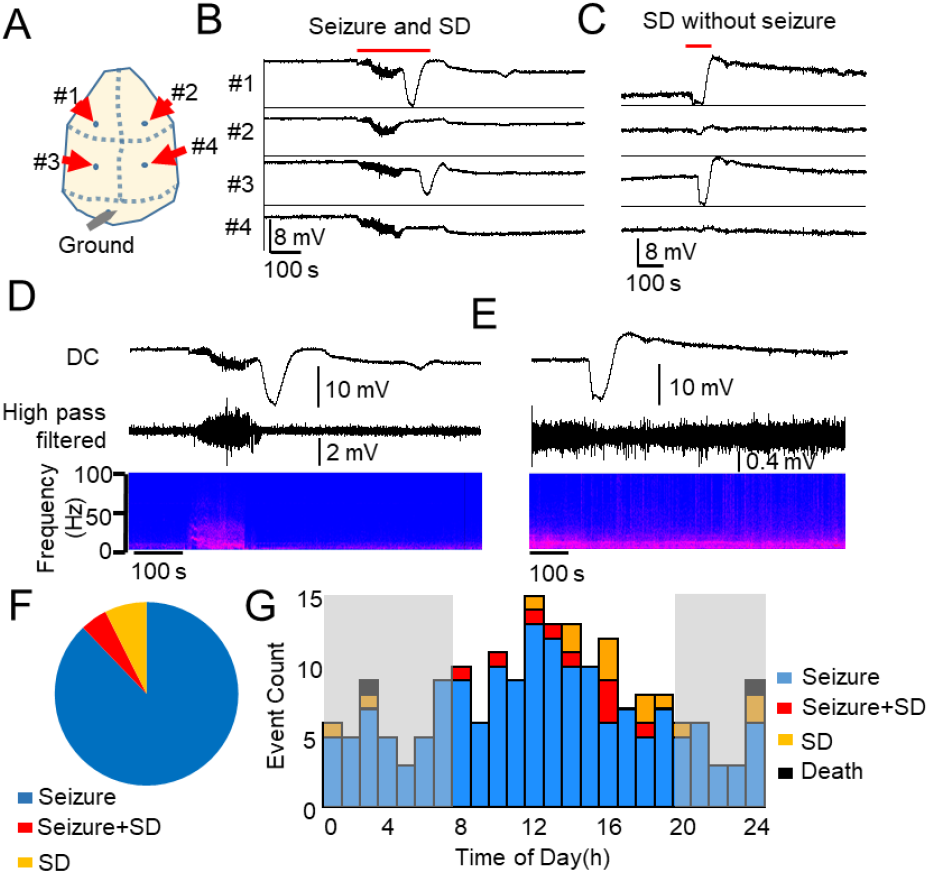
Spontaneous bilateral seizure with unilateral SD and seizure independent SD in Kcna1 KO mice. Chronic DC recording in juvenile Kv1.1/Kcna1 KO mouse. A. Representative trace of SD following a seizure (Postictal SD, B) and isolated unilateral SD (C). D&E EEG characteristics of SDs. No clear EEG suppression was detected in these recordings. F. Unlike Kcnq2-cKO mouse, seizure (n=166) and SD (n=14) were often detected independently, and a seizure+SD complex accounted for 4.8% (9/189) of total events. F. Histogram of seizure, SD and seizure+SD detected from 8 Kv1.1 KO mice. There was no clear circadian dependency.

As shown in Figure 2B&C, 74% (17/23) of spontaneous SD events in Kv1.1 KO mice were unilaterally generated even when detected following a bilateral (i.e. generalized) seizure. 13% (3/23) of the SD events were near-simultaneously detected in both hemispheres with a mean latency of 18.6 ± 1.6 s, whereas this latency is still much longer than that seen in Kcnq2-cKO mice (2.5 ± 3.6s). Unlike Kcnq2-cKO mice, in Kv1.1 KO mice, 87% (20/23) of SDs were detected first in the anterior electrodes (**Figure 2B&C**). There was slight EEG suppression following the SD (**Figure 2D&E**), which was less pronounced compared with those reported previously in anesthetized animals. Also unlike the cKo model, no clear circadian dependency was detected (**Figure 2G**).

Together these results indicate that the susceptibility of cortical SD is strongly linked to intrinsic membrane excitability characteristics regulated by Kv7.2 and Kv1.1 voltage-gated potassium channels, while their regional and temporal thresholds are largely dependent on the specific channel subtype.

### Kcnq/Kv7 channel inhibition in Kv1.1 KO mice partially phenocopies SD pattern of Kcnq2-cKO mice

The SD pattern in Kcnq2-cKO cortex is unusual in that these events are tightly linked to preceding seizures and are generated bilaterally. In order to test whether a loss of Kv7 current directly contributed to the SD generation pattern, we administered the nonselective Kv7/Kcnq inhibitor XE991 (5-10 mg/kg, i.p.) to 5 Kv1.1 KO mice and monitored spontaneous cortical EEG activity. In all Kv1.1 KO mice, bilateral seizure-SD complexes were triggered shortly after drug injection (9.9 ± 3.9 minutes). Three of these mice showed almost identical short-latency bilateral SD temporal onset, while the other 2 showed a delayed onset (33.0 and 13.3 s) between hemispheres. The calculated SD propagation rate of the SDs generated following XE991 administration was faster than the spontaneous SDs of untreated Kv1.1 KO (after XE991: 6.4 ± 2.3 mm/min, n=5, spontaneous SD: 3.2 ± 2.5 mm/min, n=20; p=0.0041 Mann Whitney U-test).

While cortical SD did not provoke noticeable motor behavior in either model, unlike the spontaneous seizure/SD complex in Kcnq2-cKO mice, the XE991-induced seizure-SD complex was lethal in Kv1.1 KO mice. We found that 3 Kv1.1 KO mice barely survived the SD occurrence and subsequently died with a latency of 5.2 ± 0.9 minutes (**Figure 3A**), and the other 2 Kv1.1 KO mice died suddenly during the repolarization phase of the DC shifts (**Figure 3A**, death was determined by the irreversible loss of spontaneous EEG activity). It is noteworthy that 3 of these Kv1.1 KO mice displayed seizures but never postictal SD during > 48-hour untreated pre-drug baseline recordings, suggesting that pharmacological Kv7 inhibition not only reproduced the bilateral SD generation pattern of Kcnq2 mutants but also facilitated postictal SD generation. No seizures or SD were detected in WT mice following XE991 administration within 24 hours of administration (n=4, **Figure 3C**), suggesting the drug had no proconvulsant or SD triggering effects.

**Figure 3.**
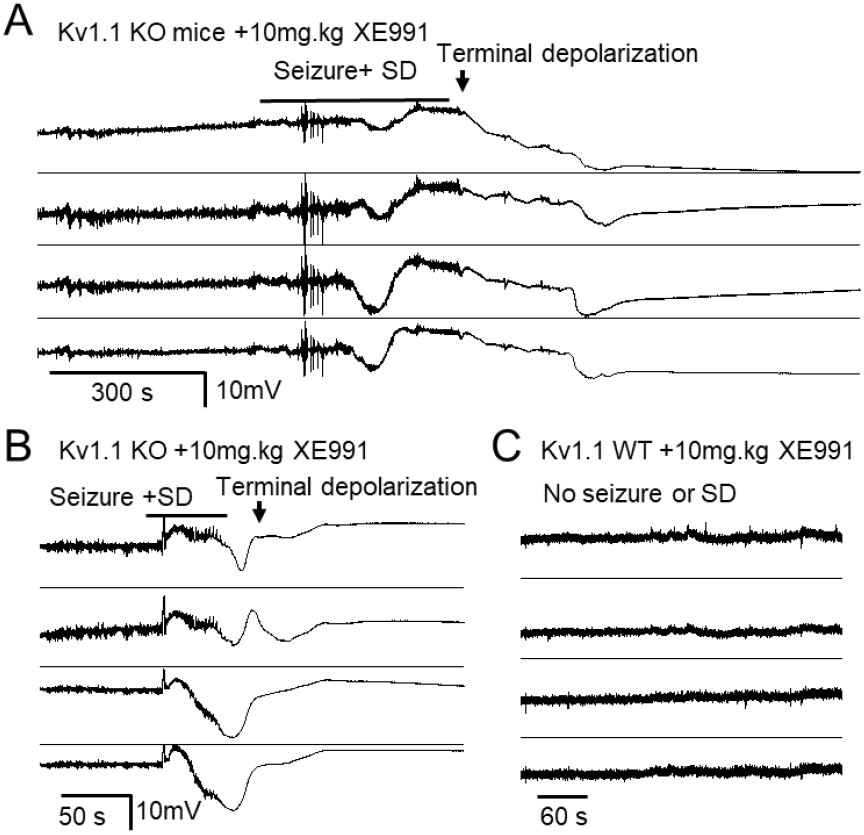
Kv7 inhibitor XE991 triggers bilateral seizure/SD complex and death in Kv1.1 KO mice. After baseline recordings, Kv1.1 KO (n=5) and WT (n=4) mice were administered a single dose of XE991 (5-10 mg/kg i.p.), triggering a bilateral seizure-SD complexes in all Kv1.1 KO mice. **A**. 60% (3/5) of KO mice acutely survived the seizure-SD complex but subsequently died. **B**. Remaining mice (2/5 mice) died immediately following the bilateral seizure-SD. No seizure or SD were detected in 4 Kv1.1 WT control mice and all survived.

Together, these DC recording studies in awake epileptic mice suggest a critical functional role of Kv7.2 current in regulating bilateral SD generation, and a distinctive SD phenotype between Kcnq2 and Kcna1 mutants.

### SD threshold is lowered in Kcnq2-cKO cortical slices *in vitro*

Since Emx1-cre and Kcnq2 are expressed in the peripheral autonomic nervous system ^32^, the prominent SD phenotype of Kcnq2-cKO mice may be attributable to changes in systemic physiological reactivity that could compromise blood flow. We therefore next examined whether the SD threshold of cortical networks is intrinsically altered in cortical brain slices.

The first set of experiments examined SD generation using a seizure induction model. Cortical slices were incubated in nominally Mg^2+^ free solution to increase spontaneous tissue excitability while monitoring local field potentials and the intrinsic optical signal (IOS). This condition produced a brief episode of spontaneously fast activity in extracellular field electrodes, often followed by SD, in both WT and mutant cortex. The latency until SD onset was significantly faster in the cKO than WT (WT: 10.6 ± 2.0 minutes, cKO: 7.7 ± 1.2 minutes, p<0.001). With this global increase in cellular excitability, we found that SDs were generated in multiple confluent foci, and we could not reliably calculate a propagation rate. These results indicate that the Kcnq2 cKO cortical slice is intrinsically more susceptible to SD generation under pro-epileptic conditions.

One mechanism underlying the higher SD susceptibility of isolated cKO cortical tissue could be attributable to altered sensitivity to extracellular K^+^. Therefore we next determined the K^+^ contribution to SD threshold under physiological conditions by incrementally elevating the bath K^+^ concentrations while monitoring pairs of cortical slices (WT and cKO) as described above. These experiments revealed a subtle but significant reduction of the K^+^ threshold in the cKO cortical slices (**Figure 4C&D;** WT: 11.2 ± 0.2 mM, cKO: 10.6 ± 0.2 mM, n=14 each, p<0.05).

**Figure 4.**
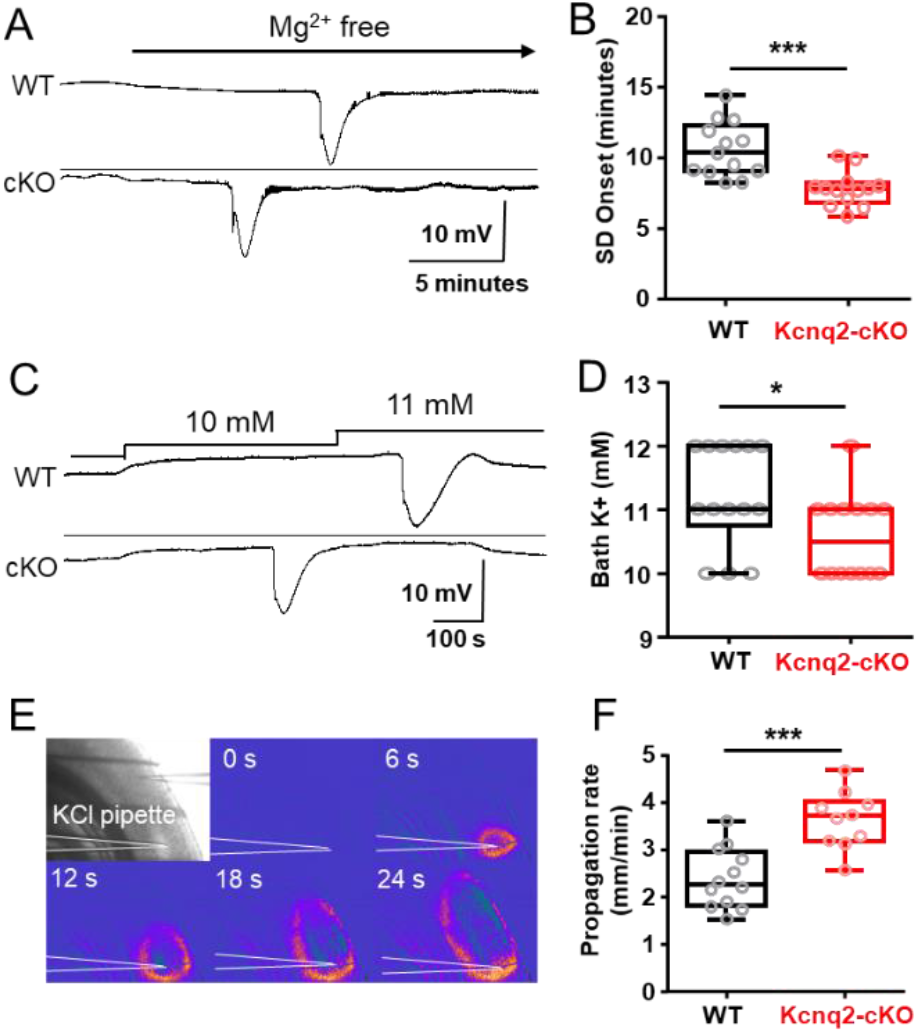
SD threshold is lowered in Kcnq2-cKO cortical slices in vitro. **A&B** SD onset following exposure to nominally Mg^2+^ free solution is faster in cKO mouse cortical slices. Representative traces (**A**) and quantification (**B**) of wild type (n=13 (WT), and mutant n=12 (cKO) slices **C&D**. Kcnq2-cKO cortical slices had a lowered K^+^ SD threshold. Pairs of slices were incubated and bath K^+^ concentration was incrementally elevated until SD is detected. (C-D) SD was evoked at lower bath K^+^ concentration in Kcnq2-cKO than WT slices (n=14 each). **E&F**. SD propagation rate was faster in Kcnq2-cKO slices. SD was evoked by focal KCl microinjection and detected by IOS (see Methods). The velocity of SD propagation was faster in the Kcnq2-cKO slices (n=12 (WT), 10 (cKO). * p<0.05, *** p<0.005

We also examined whether SD propagation is altered in cKO tissue using KCl microinjection model where, unlike the bath application model, a focally generated SD propagates through non-depolarized cortical tissue. In these experiments, SD was triggered by KCl microinjection (1M KCl, 50 psi) into layer 2/3 and the propagation rate was visually analyzed by tracking the intrinsic optical signal (IOS) in an area >500 µm away from the KCl injection site (**Figure 4E**) where the arrival of IOS coincided with the DC potential shift (**Figure 4F**). In agreement with the lower KCl SD threshold, we found that SD propagation speed was also faster in mutant cortical slices compared with the littermate control (WT: 2.4 ± 0.2 mm/min, cKO: 3.6 ± 0.2 mm/min, n=12 (WT), 10 (cKO), p<0.005).

### SD thresholds are not different between Kv1.1 WT and KO cortex *in vitro*

We measured SD susceptibility of Kv1.1 WT and KO cortical tissues prepared from P30-40 mouse pairs. Unlike Kcnq2-cKO tissue, there were no differences in SD thresholds (**Figure 5** WT: 10.58 ± 0.3 mM, cKO: 10.9 ± 0.5 mM, n=12 and 14 respectively) as determined by K^+^ bath application (Figure 5A) or the propagation rates (WT: 2.7 ± 0.1 mm/min, KO: 2.8 ± 0.1 mm/min, n=17 and 13 respectively) as determined by KCl microinjection model (**Figure 5B**).

**Figure 5.**
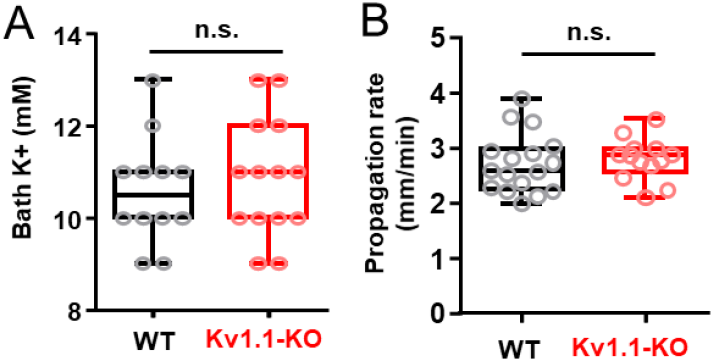
SD susceptibility is unchanged in Kv1.1 KO mouse cortical slices in vivo. **A**. SD threshold determined by incrementally increasing bath K^+^ concentrations n=12 (WT), 14 (KO). **B**. SD propagation rate was determined by focal KCl microinjection. n=17(WT), 13 (KO)

Together, these results indicate that genetic deficiency of Kv7.2, but not Kv1.1 containing channels significantly increases intrinsic susceptibility to SD. These different effects likely contribute to the high seizure-SD coupling and unique SD propagation pattern manifested *in vivo* in genetically and pharmacologically Kv7.2 depleted cortex.

### Effect of pharmacological inhibition and activation of Kv7 currents on SD in vitro

We next examined the acute modulatory activities of Kv7 channel blockade or activation on SD studied *in vitro* using XE991, a Kv7 inhibitor, and retigabine, a Kcnq2 opener. The effects on SD propagation rate were measured using the KCl microinjection model described above (**Figure 4**), in which SD was repetitively generated in WT cortical slices (by KCl puffs) while incubated with vehicle (0.1% DMSO) or XE991 (50 µM, a dose expected to fully inhibit Kv7.2 containing channels ^40^) followed by wash-out. XE991 did not modulate SD propagation rate (Control: 2.4 ± 0.2 mm/min, XE991: 2.8 ± 0.5 mm/min, Wash: 2.6 ± 0.6 mm/min, n=6, p=0.37, **Figure 6A**). In contrast, XE991 slightly reduced the KCl threshold when examined using the KCl bath application model (Control: 11.0 ± 0.2 mM, XE991: 10.0 ± 0.2 mM, n=12, p<0.005, **Figure 6B**). Taken together, Kv7 inhibitor has a marginal facilitatory effect on SD generation *in vitro*.

**Figure 6.**
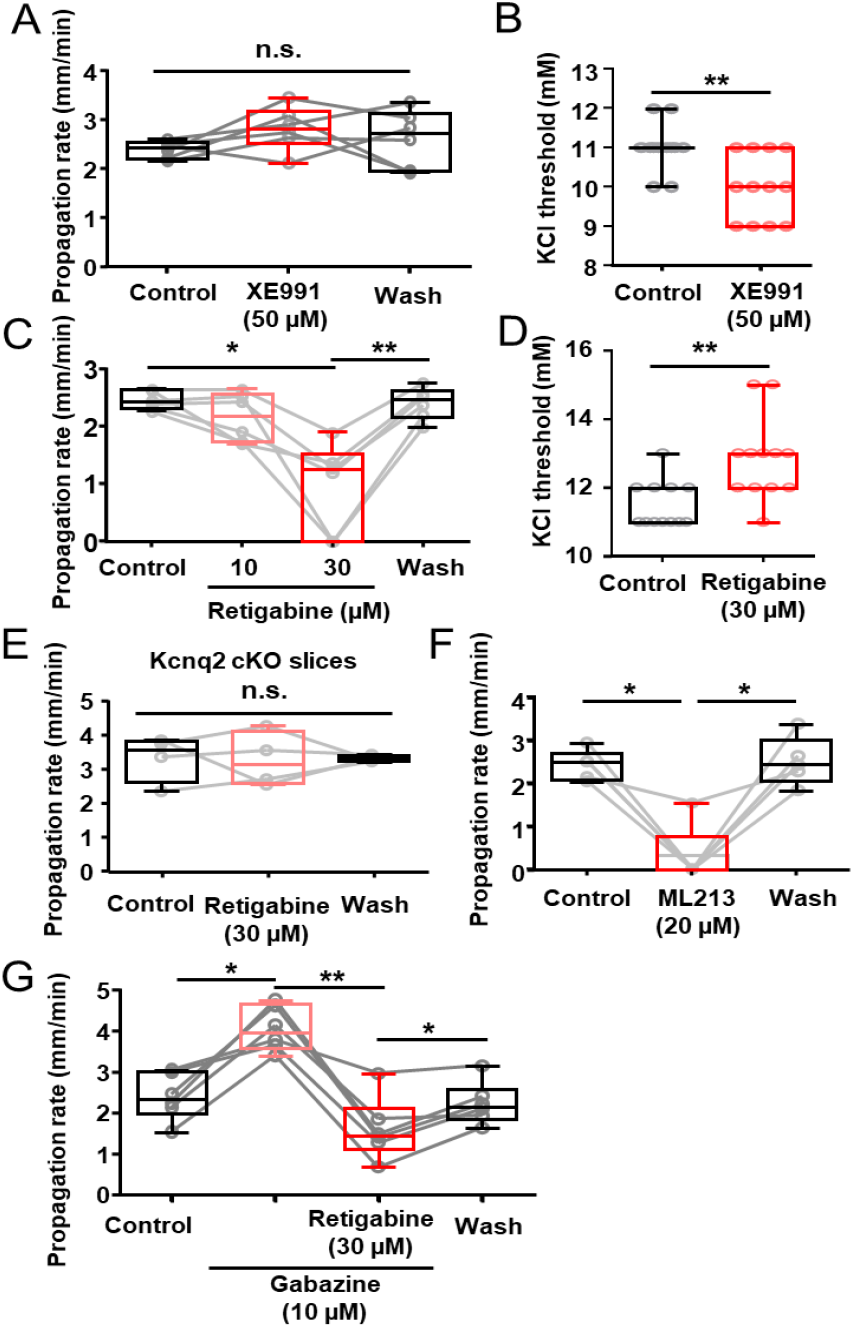
Acute Enhancement/Inhibition of SD in WT cortex by Kv7 inhibitor/activator in vitro. **A&B**. Effect of XE991 on SD propagation rate and K^+^ threshold. **A**. SD propagation rate measured from SD repetitively generated in single slices while incubated in the drug was unchanged. n.s = not significant, n=6. **B**. SD threshold was decreased by Kcnq inhibitor in K^+^ bath application model. n=12 each * p< 0.05. **C&D**. Retigabine dose-dependently reduced SD propagation rate (**C**, n=6) and elevated the K^+^ threshold (**D**, n=12). **E**. Retigabine (30 µM) did not alter SD propagation in Kcnq2-cKO slices. n=4 **F**. ML216 (20 µM), another Kv7 activator, also inhibited SD propagation in WT slices. n=5 **G**. Retigabine effect on SD was analyzed in the presence of the GABAAR antagonist gabazine. Gabazine increased the SD propagation rate, however

We next examined the effect of retigabine, a clinically used non-selective Kv7 current activator (^41^, **Figure 6B&C**). Retigabine was without effect at 10 µM, while at 30 µM, retigabine either completely blocked SD (2/6) or greatly reduced the speed of propagation (4/6 slices). This inhibitory effect was fully reversed by extensive drug washout (> 30 min). Retigabine also elevated the KCl threshold of SD as measured in the incremental KCl bath application model (vehicle: 11.5 ± 0.2 mM retigabine: 12.8 ± 0.3 mM, respectively p=0.016, **Figure 6D**).

The retigabine effect is largely due to the modulation of the Kv7 channels since the inhibitory effect was absent when tested on Kcnq2-cKO brain slices (**Figure 6E**). ML216, a structural analog of retigabine and more selective Kv7.2 and 7.4 channel opener ^42(p213)^, also fully inhibited SD generation in 80% (4/5) of experiments and reduced SD propagation in the rest (Control: 2.4 ± 0.4 mm/min, ML216: 0.3 ± 0.7 mm/min, wash: 2.5 ± 0.6 mm/min, n=5 **Figure 6F**).

In addition to Kv7 channels, retigabine has an off-target potentiating effect on GABA_A_R currents ^43,44^. We therefore examined the GABA_A_R dependency of retigabine using repetitive KCl evoked SD in acute cortical slices. Consistent with previous studies, exposure to the GABA_A_R antagonist, gabazine (10µM) significantly increased the SD propagation rate (Control: 2.4 ± 0.6 mm/min, gabazine: 4.1 ± 0.5 mm/min). Retigabine was still effective in this condition, consistently reducing SD velocity (gabazine + retigabine: 1.6 ± 0.8 mm/min), and the effect was reversible (washout retest velocity: 2.2 ± 0.5 mm/min)). These results indicate that SD inhibition by retigabine is independent of GABA_A_R signaling (**Figure 6G**). The SD inhibitory effect of retigabine was further tested in the mouse cortex *in vivo*. In urethane anesthetized WT adult mice, SD was elicited by superfusing incrementally increasing concentrations of KCl (100, 250, 500, 1000 mM, 30 s) onto the pial surface through a cranial window created over the posterior cortex and detected with both a surface electrode as well as transcranial imaging of the intrinsic optical signal (light scatter, **Figure 7A)**. In each animal, the SD threshold was determined twice, first at pre-drug baseline, and then 20 minutes after i.p injection of drug or DMSO vehicle control. In these experiments, 30 mg/kg retigabine (but not 10 mg/kg) significantly increased the KCl threshold of SD required to trigger an SD (**Figure 7B&C**).

**Figure 7.**
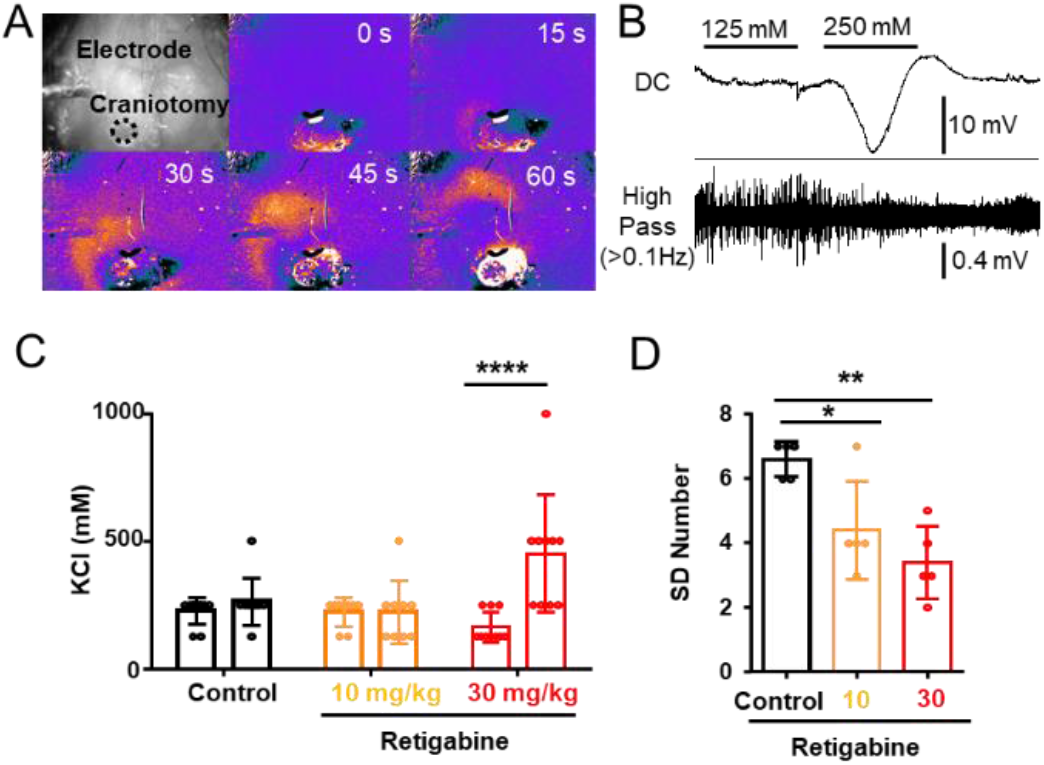
SD inhibition by retigabine in in vivo anesthetized WT mouse cortex. SD was triggered by repetitively applying a KCl solution (100, 300, 500, 1000 mM for 2 minutes) to the cortical surface. A. SD wave was detected with IOS signal shown in a pseudocolored image, and B. electrophysiologically with Ag/AgCl electrode. **C**. Summary of SD threshold measurement. In each animal, SD threshold was measured before and after drug injection. Vehicle (0.1% DMSO) and 10 mg/kg retigabine had no effect, while 30 mg/kg retigabine significantly increased K^+^ evoked SD threshold. ****p<0.001 **D**. Number of recurrent SD events during continuous 0.5 M KCl application for 30 minutes. Retigabine 30mg/kg significantly decreased regenerative SD number. *p<0.05, ** p<0.01

After testing the SD threshold, the frequency of recurrent SD generation during a continuous 0.5M KCl application was determined in a subset of animals. Consistent with an elevated SD threshold, the number of recurrent SDs in response to a continuous surface application of 0.5M KCl was decreased following retigabine treatment (**Figure 7D**).

Together these results indicate that Kv7 inhibition and activation have modulatory effects on SD in the WT brain. Activation seems to have a stronger effect on threshold than inhibition (see Discussion). Given the pharmacological concentration range of retigabine on Kcnq current (0.1 – 100 µM ^45^), these data also suggest that SD inhibition required near-maximal concentrations for an anti-SD effect.

### Retigabine inhibits SD in mild, but not severe oxygen-glucose deprivation (OGD) model

Kv7 activators have been shown to reduce neuronal damage following experimental ischemia, where SD could potentially contribute to progression of the lesion ^46,47^. We examined whether the Kv7 activator retigabine can also inhibit SD generated during the experimental in vitro ischemia model of oxygen-glucose deprivation (OGD). In this model, exposure to OGD solution (0% O_2_, 0 or 2 mM glucose, see Methods for detail) results in spontaneous SD generation (**Figure 8A&B**), and the latency to the first SD after OGD exposure is used as a readout of SD threshold. In this model, cortical slices were resistant to SD inhibition by retigabine (30 µM) when exposed to a 0% O_2_/0 mM glucose solution (**Figure 8C**). However, retigabine significantly delayed SD onset when bathed with 0% O_2_/2 mM glucose OGD solution (**Figure 8C**). Thus the acute inhibitory effect of retigabine appears limited under conditions of mild hypoxic/metabolic deprivation, but not effective when glucose metabolism is severely compromised.

**Figure 8.**
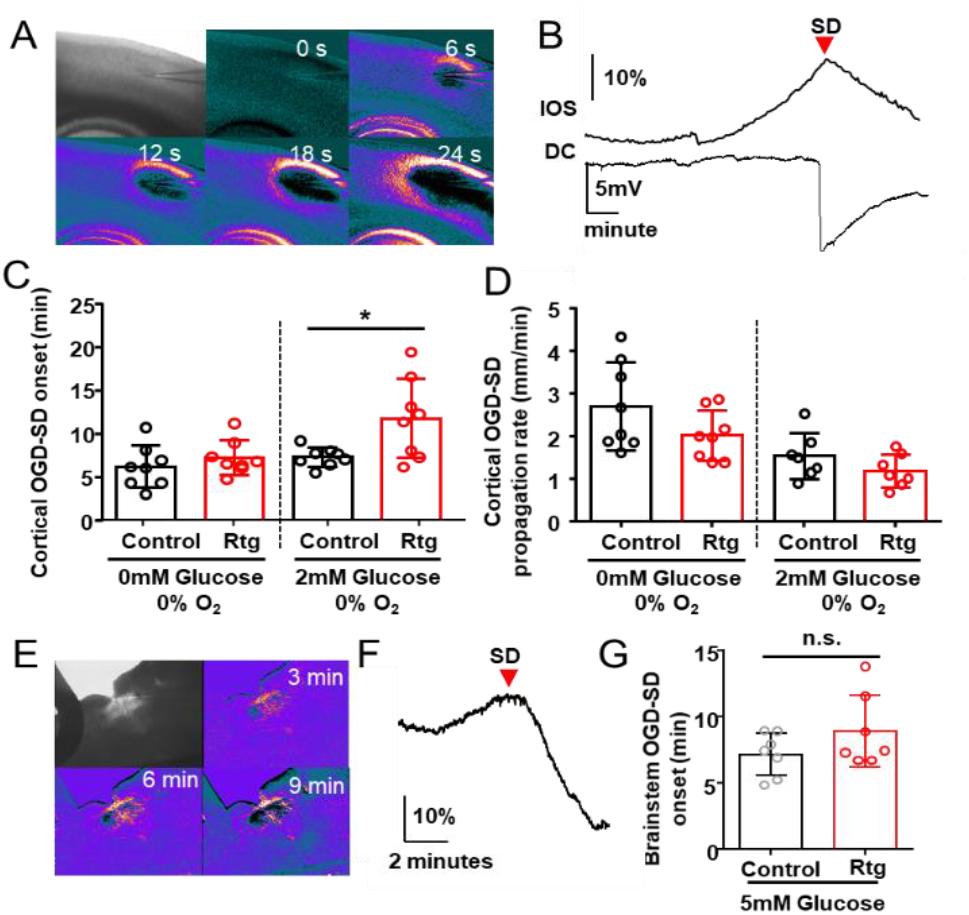
A-D Kv7.2 activator retigabine inhibits mild, but not severe OGD-induced SD. **A**. Image of OGD-SD wave in cortical slice triggered by continuous exposure to OGD solution (0% O2, 0 mM glucose), and detected as IOS traveling across cortical tissue. **B**. Arrival of IOS signal (upper) near electrode coincided with negative DC potential shift (lower trace). **C&D**. Retigabine delayed OGD-SD when metabolic stress was mild (2mM glucose), but had no effect during severe compromise (0mM glucose). Retigabine did not inhibit SD propagation rate (**D**). n=8 each, * p<0.05 **E-G**. Retigabine had no effect on OGD-SD generated in slices of the medulla at the level of the nucleus tractus solitarius (nTS). **E**. SD was triggered by 0%O_2_/5 mM glucose solution with or without 30 µM retigabine. **F, G**. SD onset was determined by the IOS signal peak at the lateral

To determine whether retigabine might be effective in subcortical brain regions where seizure-induced SD has been linked to local hypoxia and cardiorespiratory arrest ^3^, we examined its effect on SD evoked by OGD in slices of medulla containing the nucleus tractus solitaries (nTS). Some ion channel mutations implicated in SUDEP, including loss of Kv1.1, significantly lower SD threshold in this region ^16,48^. The effect of retigabine was tested in brainstem slices prepared from juvenile Kv1.1 KO mice and based on the latency to OGD-SD onset determined by the IOS changes (**Figure 8E-G**). Retigabine did not block or delay SD onset in this model under 5 mM glucose OGD conditions (**Figure 8G**). Thus the inhibitory effect of retigabine on brainstem SD in vivo is model-dependent and likely influenced by both the severity of metabolic stress and the functional contribution of Kcnq2 channels in different anatomical networks.

## Discussion

Our study identifies a novel role of Kcnq2/Kv7.2 in the regulation of susceptibility to pathological long-lasting depolarization of cortical networks. Chronic DC recordings in mice as early as 24 days old revealed a distinctive cortical SD phenotype when Kv7.2 subunits are genetically removed or pharmacologically inhibited compared to a related axonal potassium channel, Kcna1/Kv1.1. Unlike Kv1.1 deficient mice, loss of Kv7.2 in excitatory neurons facilitated tightly coupled seizure-SD transitions, bihemispheric SD generation, and a circadian expression pattern. *In vitro* analyses of acute cortical slices in response to multiple SD triggers revealed that the lower SD threshold is intrinsic to cortical tissue excitability and that Kv7.2 activity bidirectionally regulates K^+^ SD thresholds. These results support Kcnq2/Kv7.2 availability as a molecular determinant of SD threshold and cortical distribution. Fluctuations of this current under pathological states ^49–51^ may contribute to SD susceptibility, and serve as a therapeutic target in epilepsy and other neurological disorders

### Kcnq2/Kv7.2 regulates seizure/SD transition

Seizures in Kcnq2 mice reliably triggered SD expression 97% (34/35 events) of the time, and at very short latency, sometimes even commencing during the seizure. This is in sharp contrast to Kv1.1 KO mice, in which only 5% (9/175) of seizures were associated with SD, with a significantly longer postictal latency. The severity of the seizure itself is unlikely to explain these differences, as seizure duration was longer in Kv1.1 KO mice and should create an even larger extracellular potassium surge and tissue hypoxia. Instead, the conversion of uncoupled seizure/SD to a tightly coupled Kcna2-cKO-like SD phenotype by acute pharmacological inhibition (**Figure 3**) indicates the presence of a functional Kcnq2/Kv7.2 activity-dependent threshold mechanism. While the threshold is multifactorial and state-dependent, our study suggests that reducing Kv7.2 current lowers the K^+^ threshold for SD generation during a seizure. In addition to the SD threshold, changes in tissue metabolism, astrocytic extracellular homeostasis, and synaptic network functions during chronic epileptic conditions may also modulate the seizure-SD transition *in vivo* ^52,53^.

Although they were rare, the presence of seizure-independent isolated SD events in Kv1.1 KO mice was somewhat unexpected (**Figure 2C**). The triggering mechanism of these isolated events is not known, but may reflect focal changes in cortical hyperexcitation or vascular insufficiency, since cerebral autoregulation may be impaired in some forms of epilepsy ^54,55^.

### Kcnq2/Kv7.2 regulation of bihemispheric SD generation

Another salient feature of reduced Kv7.2 availability is the bihemispheric symmetry of SD following a generalized seizure. The majority of SD-related neurological deficits during migraine auras such as visual scotomata and hemiplegia are unilateral ^56^. Short latency, bihemispheric SD generation in this model is likely a direct consequence of the preceding generalized seizure activity, which raises extracellular K^+^ and lowers tissue oxygenation in both hemispheres simultaneously. A similar bilateral SD generation pattern has been reported following a generalized seizure induced by systemic chemoconvulsant (PTZ) injection ^15^. In this regard, the predominant unilateral SD in Kv1.1 KO mice after a bilateral generalized seizure seems more difficult to explain (**Figure 2B**), however the fact that acute Kv7 inhibition produced bihemispheric SD generation in Kv1.1 KO mice indicates that Kv7 activity acts as a barrier for bilateral generation. Based on its enrichment within axonal initial segments and nodes ^25,33^, Kv7.2 activity may limit long-range (e.g. transcallosal) axonal propagation of hyperactivity, thereby reducing the likelihood of contralateral SD generation in symmetric brain regions. This mechanism might also contribute to the faster tissue propagation velocity into the surrounding neocortex by activation of shorter range subcortical association pathways.

### SD modulation by Kv7.2 activity

Kcnq2/Kv7.2 containing channels generate a non-inactivating outward current that plays a critical role in limiting the generation of persistent sodium current (INaP, ^57^) an initial inward current of SD ^1^.

Genetic deletion or pharmacological inhibition of M-current likely facilitates INaP and accelerates the membrane depolarization, while augmentation may attenuate INaP, which may elevate the SD threshold. This relationship is in contrast to Kv1.1, which acts to limit action potential bandwidth and frequency ^38 25,58^. Loss of this fast current may be critical for promoting synchronous action potential bursting leading to seizures, but might be less important as a brake for slow sustained cellular depolarization leading to SD. In addition to a direct genetic effect, biological changes during chronic epilepsy might also affect SD thresholds, and we have observed elevated cortical SD threshold in older Kv1.1 KO mice.

We showed that retigabine could delay SD onset following submaximal OGD stimulation. Interestingly, Kv7.2 activators are neuroprotective in experimental ischemia and brain trauma studies ^46,47,51,59,60^, and the anti-SD properties of the activator may contribute to these neuroprotective effects. It should be noted that only a relatively high, sedating, drug concentration effectively inhibited SD, and the drug may therefore be of limited prophylactic utility. On the other hand, Kv7 activators might find an adjunctive role in sedated patients requiring critical care. We also found asymmetric effects of the Kcnq2 activator and inhibitor. The Kv7 inhibitor had a relatively small facilitatory effect on SD generation, whereas the activator could fully suppress SD at a high concentration. This result may indicate relatively small basal Kv7 current amplitudes, with a capacity to generate larger currents upon potentiation by GPCR signaling and binding of endogenous agonists ^61,62^. It has also been suggested that the Kv7 activator retigabine may indirectly antagonize NMDARs, which are important contributors to SD initiation/propagation ^63^. This mechanism could also contribute to SD inhibition by Kv7 activators (**Figure 4-6**), however *in vivo* evidence of functional NMDAR inhibition by retigabine is limited and requires further study.

### Circadian dependency of seizures in potassium channel epileptic mice

Finally, we identified a distinctly different circadian pattern of spontaneous SD generation between the Kv1.1 KO and Kcnq2-cKO potassium channelopathy models (**Figure 1&2**). In both models, we detected spontaneous seizures and SD, however a preponderance of seizure-SD complexes in the Kcnq2-cKO model occurred during the dark phase when mice are active. Strong diurnal periodicity of excitability and behavior is well known in genetic epilepsies ^64^, and several potassium channels expressed in suprachiasmatic pathways have been recently implicated in this process ^65–67^. These channels may contribute to the circadian independent seizure expression in Kv1.1 KO mouse.

## Acknowledgments

We thank Dr A. Tzingounis for generously sharing Kcnq2-flox mice.

## Figure legends

**Figure 1. Spontaneous Seizure/SD complexes predominate in Emx1-Kcnq2cKO mice**

Chronic cortical DC recording from juvenile Kcnq2-cKO mouse. A. Recording configuration. B&C. Spontaneous seizure/SD complex in awake Kcnq2-cKO mouse shown at slow (B) and fast (C) time scales. SD was detected as a sharp negative DC potential shift following a brief generalized seizure detected simultaneously in both hemispheres. A solitary ectopic DC shift was occasionally detected posteriorly (C, arrowhead). D. characterization of fast EEG activity during seizure and SD. The negative DC shifts of SD were detected immediately after or even before the end of seizure activity. Left: slow time scale, right: faster scale. High-frequency EEG activity was enhanced during the nadir of the DC shift (b), followed by suppression with occasional spikelets (c). E. Almost all (97%, n= 34/35) spontaneous seizures were followed by SD. No SD’s were detected without a preceding seizure. F. Circadian cycle of seizure/SD complexes in 7 Kcnq2-cKO mice. Histogram of seizure and SD incidence during 24 h period (bin size = 1 hour, dark gray = dark phase). Spontaneous seizure/SD complexes were detected primarily during the dark phase (p<0.0001, Fisher’s exact test).

**Figure 2. Spontaneous bilateral seizure with unilateral SD and seizure independent SD in Kcna1 KO mice.**

Chronic DC recording in juvenile Kv1.1/Kcna1 KO mouse. A. Representative trace of SD following a seizure (Postictal SD, B) and isolated unilateral SD (C). D&E EEG characteristics of SDs. No clear EEG suppression was detected in these recordings. F. Unlike Kcnq2-cKO mouse, seizure (n=166) and SD (n=14) were often detected independently, and a seizure+SD complex accounted for 4.8% (9/189) of total events. F. Histogram of seizure, SD and seizure+SD detected from 8 Kv1.1 KO mice. There was no clear circadian dependency.

**Figure 3. Kv7 inhibitor XE991 triggers bilateral seizure/SD complex and death in Kv1.1 KO mice.**

After baseline recordings, Kv1.1 KO (n=5) and WT (n=4) mice were administered a single dose of XE991 (5-10 mg/kg i.p.), triggering a bilateral seizure-SD complexes in all Kv1.1 KO mice. **A**. 60% (3/5) of KO mice acutely survived the seizure-SD complex but subsequently died. **B**. Remaining mice (2/5 mice) died immediately following the bilateral seizure-SD. No seizure or SD were detected in 4 Kv1.1 WT control mice and all survived.

**Figure 4. SD threshold is lowered in Kcnq2-cKO cortical slices in vitro.**

**A&B** SD onset following exposure to nominally Mg^2+^ free solution is faster in cKO mouse cortical slices. Representative traces (**A**) and quantification (**B**) of wild type (n=13 (WT), and mutant n=12 (cKO) slices **C&D**. Kcnq2-cKO cortical slices had a lowered K^+^ SD threshold. Pairs of slices were incubated and bath K^+^ concentration was incrementally elevated until SD is detected. (C-D) SD was evoked at lower bath K^+^ concentration in Kcnq2-cKO than WT slices (n=14 each). **E&F**. SD propagation rate was faster in Kcnq2-cKO slices. SD was evoked by focal KCl microinjection and detected by IOS (see Methods). The velocity of SD propagation was faster in the Kcnq2-cKO slices (n=12 (WT), 10 (cKO). * p<0.05, *** p<0.005

**Figure 5. SD susceptibility is unchanged in Kv1.1 KO mouse cortical slices in vivo.**

**A**. SD threshold determined by incrementally increasing bath K^+^ concentrations n=12 (WT), 14 (KO). **B**. SD propagation rate was determined by focal KCl microinjection. n=17(WT), 13 (KO)

**Figure 6. Acute Enhancement/Inhibition of SD in WT cortex by Kv7 inhibitor/activator in vitro.**

**A&B**. Effect of XE991 on SD propagation rate and K^+^ threshold. **A**. SD propagation rate measured from SD repetitively generated in single slices while incubated in the drug was unchanged. n.s = not significant, n=6. **B**. SD threshold was decreased by Kcnq inhibitor in K^+^ bath application model. n=12 each * p< 0.05. **C&D**. Retigabine dose-dependently reduced SD propagation rate (**C**, n=6) and elevated the K^+^ threshold (**D**, n=12). **E**. Retigabine (30 µM) did not alter SD propagation in Kcnq2-cKO slices. n=4 **F**. ML216 (20 µM), another Kv7 activator, also inhibited SD propagation in WT slices. n=5 **G**. Retigabine effect on SD was analyzed in the presence of the GABA_A_R antagonist gabazine. Gabazine increased the SD propagation rate, however retigabine still reduced the SD propagation rate. * p<0.05, ** p<0.01

**Figure 7. SD inhibition by retigabine in in vivo anesthetized WT mouse cortex.**

SD was triggered by repetitively applying a KCl solution (100, 300, 500, 1000 mM for 2 minutes) to the cortical surface. A. SD wave was detected with IOS signal shown in a pseudocolored image, and B. electrophysiologically with Ag/AgCl electrode. **C**. Summary of SD threshold measurement. In each animal, SD threshold was measured before and after drug injection. Vehicle (0.1% DMSO) and 10 mg/kg retigabine had no effect, while 30 mg/kg retigabine significantly increased K^+^ evoked SD threshold. ****p<0.001 **D**. Number of recurrent SD events during continuous 0.5 M KCl application for 30 minutes. Retigabine 30mg/kg significantly decreased regenerative SD number. *p p<0.05, ** p<0.01

**Figure 8**

A-D Kv7.2 activator retigabine inhibits mild, but not severe OGD-induced SD. **A**. Image of OGD-SD wave in cortical slice triggered by continuous exposure to OGD solution (0% O2, 0 mM glucose), and detected as IOS traveling across cortical tissue. **B**. Arrival of IOS signal (upper) near electrode coincided with negative DC potential shift (lower trace). **C&D**. Retigabine delayed OGD-SD when metabolic stress was mild (2mM glucose), but had no effect during severe compromise (0mM glucose). Retigabine did not inhibit SD propagation rate (**D**). n=8 each, * p<0.05 **E-G**. Retigabine had no effect on OGD-SD generated in slices of the medulla at the level of the nucleus tractus solitarius (nTS). **E**. SD was triggered by 0%O_2_/5 mM glucose solution with or without 30 µM retigabine. **F, G**. SD onset was determined by the IOS signal peak at the lateral margin of the nTS. (n=7 in all experiments).

